# HtrA1 activation is driven by an allosteric mechanism of inter-monomer communication

**DOI:** 10.1101/163717

**Authors:** Alvaro Cortes Cabrera, Esther Melo, Doris Roth, Andreas Topp, Frederic Delobel, Corinne Stucki, Chia-yi Chen, Peter Jakob, Balazs Banfai, Tom Dunkley, Oliver Schilling, Sylwia Huber, Roberto Iacone, Paula Petrone

## Abstract

The human protease family HtrA is responsible for preventing protein misfolding and mislocalization, and a key player in several cellular processes. Among these, HtrA1 is implicated in several cancers, cerebrovascular disease and age-related macular degeneration. HtrA1 activation, although very relevant for drug-targeting this protease, remains poorly characterized. Our work provides a mechanistic step-by-step description of HtrA1 activation and regulation. We report that the HtrA1 trimer is regulated by an allosteric mechanism by which monomers relay the activation signal to each other, in a PDZ-domain independent fashion. Notably, we show that inhibitor binding is precluded if HtrA1 monomers cannot communicate with each other. Our study establishes how HtrA1 oligomerization plays a fundamental role in proteolytic activity. Moreover, it offers a structural explanation for HtrA1-defective pathologies as well as mechanistic insights into the degradation of complex extracellular fibrils such as tubulin, amyloid beta and tau that belong to the repertoire of HtrA1.

**Highlights:**

- Monomeric HtrA1 is activated by a gating mechanism.
- Trimeric HtrA1 is regulated by PDZ-independent allosteric monomer cross-talk.
- HtrA1 oligomerization is key for proteolytic activity.
- Substrate-binding is precluded if monomers cannot communicate with each other.

## INTRODUCTION

The high-temperature requirement A (HtrA) family of serine proteases prevent cellular malfunction arising from protein misfolding and mislocalization^1,2^. Human HtrA1 is responsible for the cleavage of substrates such as the tau protein and tubulin^3,4^. HtrA1 up- or down-regulation have been associated with important pathological processes. For example, a polymorphism in the promoter sequence of the HtrA1 gene results in an abnormal increase in HtrA1 protease levels, linked to age-related macular degeneration (AMD)^5^. Diseases such as the CARASIL (Cerebral Arteriopathy, Autosomal Recessive with Subcortical Infarcts and Leukoencephaolopathy) syndrome and the cerebral small vessels disease (CSVD) are caused by mutations or deletions that impair HtrA1 function^6^. HtrA1 also plays a role in tumor suppression^7^. While it stands as a promising pharmaceutical target, HtrA1 activation and regulation are not fully understood.

In solution, HtrA1 is most abundant as a trimer. Each monomer is composed of four different domains: a trypsin-like catalytic domain, a C-terminal PDZ domain, a IGFBP-like domain and Kazal-like domain^8^. In bacterial homologs DegP and DegS, the PDZ domain orchestrates activation by driving conformational change across the monomers of the oligomeric forms, upon binding of unassembled outer-membrane proteins (OMPs) accumulated in the periplasm^9,10^. However, notably in humans, the PDZ domain has proven largely dispensable for the regulation of proteolytic activity, although important for correctly processing polymeric substrates^11^. This indicates that human HtrA1 might undertake an alternative PDZ-independent activation process which is currently unknown.

Molecular dynamics (MD) simulations have been successfully applied to characterize molecular states leading to conformational change^12-14^. We combined this tool with community analysis^15^ to identify allosteric regions and to investigate the mechanisms leading to HtrA1 activation both at the monomer level and globally for the trimer.

Our aim is to understand the structural and dynamic mechanisms leading to HtrA1 activation, relevant for the development of new therapeutic strategies. We find that allostery is a driver of human HtrA1 regulation and explain how the proteolytic activation in human HtrA1 is achieved independently of the PDZ domain, mediated by inter-monomer communication. Combining computational and experimental evidence from Retinal Pigmented Epithelium (RPE cells), an AMD disease relevant model, we elucidate how an allosteric signal, transmitted throughout the amino acid network, could promote HtrA1 catalytic activity. Our study provides structural explanation for HtrA1-related pathologies such as CARASIL and CSVD, and explains why oligomeric architecture is critical to HtrA1 function.

## RESULTS

### HtrA1 monomer is activated by a gating mechanism

Molecular dynamics simulations of static X-ray structures were used to generate hypotheses on the intermediate conformational states leading to HtrA1 activation, which we later validated *in vitro* with site-directed mutagenesis.

Starting from the fully active and fully inactive HtrA1 catalytic domain conformations, a five-state hidden Markov model has been constructed making use of extensive MD conformational sampling to investigate the cascade of events leading from the inactive to the active conformations.

The protein flows through several intermediates in which the catalytic triad (Ser 328, His220 and Asp250), the L2 loop residues Leu345 and Lys346, the LD loop and the L3 loop (residues 298-307) suffer a major rearrangement to produce an active conformation (see Figure 1 for a detailed description). Hydrophobic Leu345 stands obstructing the entrance of the substrate to the binding pocket. The dynamics of the L2 loop, and particularly Lys346 are key to initiate the conformational transition to remove Leu345, in a PDZ-independent manner.

**Figure 1.**
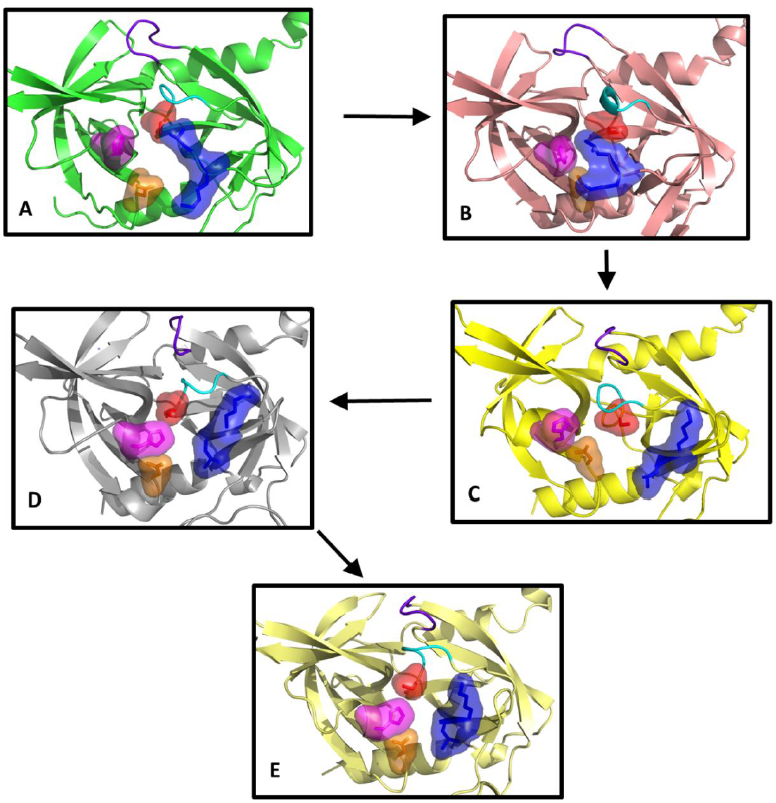
Sequence of states in the activation process from computational simulation. The residues Ser328 (surface, red), His220 (surface, magenta), Asp250 (surface, orange), L2 loop residues Leu345 and Lys346 (surface, blue), the LD loop (purple) and the oxyanion hole forming loop (cyan) are highlighted. (A) Fully inactive state that closely resembles the X-ray inactive structure (PDB 3NUM). The misaligned configuration of the catalytic triad (Ser328, His220 and Asp250) is incompatible with catalytic activity. In the binding pocket, the oxyanion hole-forming loop is disorganized and not functional while residue Leu345 occupies the cavity as a gate occluding the entrance. (B) The L2 loop adopts an intermediate conformation, where the side chain of the residues Leu345 and Lys346 are not opposed but parallel, while the rest of the elements involved in the catalytic mechanism remains similar to the inactive state A. (C) Leu345 and Lys346 have evolved to an active position from the disorganized intermediate B, inverting the orientation of their side chains originally displayed in their active conformations. This movement unblocks the S1 site of the protease while the remaining elements (catalytic triad, oxyanion hole forming loop and L3 loop) are in an inactive-like state as in state A and B. (D) The catalytic triad and all the remaining elements are now aligned in an active configuration. Only the oxyanion hole-forming loop remains disordered and not functional. (E) The structure is completely active and resembles the crystallographic active conformation (PDB 3NZI). Loop L3 stabilizes to adopt a well-defined secondary structure.

We propose a model of lock-and-gate activation by which, independently of the PDZ domain, the dynamics of the L2 loop, and particularly Lys346 (lock), would be key to initiate the transition by forcing a conformational change on Leu345, which acts as a hydrophobic gate obstructing the entrance of the substrate to the binding pocket in the inactive conformation. A similar mechanism, with tryptophan residues, has been described in other members of the trypsin-like serine protease family^16^.

The withdrawal of the Leu345 side chain from the S1 pocket into the solvent seems thermodynamically unfavorable, and therefore unlikely to happen unless a protein conformational change facilitates the process. On the other hand, the positively charged residue Lys346 is found exposed to the solvent, at different degrees, in both the active and inactive states. Our simulations suggest that the flexibility and dynamics of this side chain could potentially trigger the flip of the L2 loop that unblocks the S1 site facilitating the transition towards the active conformation.

To test this hypothesis, we first determined the kinetic parameters for the wild-type protein and then we simulated and subsequently expressed the variants L345G and K346I for biochemical enzymatic characterization (Table 1). Noteworthy, the assayed wild-type catalytic efficiency (Table 1) is in line with other serine proteases^17^, which suggests a low-energy barrier leading to HtrA1 substrate-driven activation.

**Table 1.** Kinetic parameters for wild-type HtrA1 and variants. (Supp. Figure S1, reaction rate plots).^a^ The protein is essentially inactive.

| HtrA1 variant | $K_{cat}$ ($\text{min}^{-1}$) | $K_m$ ($\mu\text{M}$) | $K_{cat}/K_m$ ($\text{M}^{-1}\text{s}^{-1}$) |
| --- | --- | --- | --- |
| Wild-type (positive control) | 14.60+-0.53 | 0.48 +- 0.03 | $5.07 \times 10^5$ |
| S328A (negative control) | Inactive<sup>a</sup> | Inactive<sup>a</sup> | Inactive<sup>a</sup> |
| R302A | Inactive<sup>a</sup> | Inactive<sup>a</sup> | Inactive<sup>a</sup> |
| E306A and R310A | 5.99 +- 0.16 | 0.28 +- 0.01 | $3.56 \times 10^5$ |
| K346I | 1.46+-0.15 | 0.67 +- 0.31 | $0.36 \times 10^5$ |
| L345G | 1.15+-0.03 | 0.14+- 0.02 | $1.37 \times 10^5$ |

Mutation K346I was designed to reduce the flexibility of Lys346 by replacing the positively charged side chain with a more conformationally restrained residue of similar size and hydrophobic character. Simulations of the K346I variant showed a very stable L2 loop, but with a looser oxyanion-forming (residues 325–327) loop due to the missing interactions of the original lysine residue. To further investigate the role of this mutation, we established a sensitive enzymatic assay for the variant using the fluorescence-quenched peptide substrate H2-Opt-previously reported as an excellent substrate for both HtrA1 and HTRA2^8^ (Methods). The biochemical characterization of K346I reported a 10-fold decrease in K_cat_ (Table 1), consistent with the proposed dynamic character of this residue and its role as initiator of the activation mechanism.

With the L345G variant, we aimed to eliminate the side chain that blocks the active site hoping to obtain an increase in activity. However, glycine substitutions are known to increase loop flexibility by removing steric interference^18.^ Simulations with the glycine variant predicted an enzyme with a large increase in L2 loop flexibility that could potentially interfere with catalysis by perturbing the configuration of the substrate and the catalytic triad. Accordingly and as expected from the flexibility shown in the simulation, this variant has greatly impaired catalytic activity (10-fold decrease in K_cat_, Table 1), highlighting the essential role of this residue (both size and dynamics) in the activity and regulation of the protease.

### Allosteric activation of HtrA1 trimer

We used molecular dynamics simulations to identify concerted motions in HtrA1 that could possibly lead to the activation process. Importantly, the pattern and distribution of dynamic groups varies between the fully inactive and fully active trimeric HtrA1. In particular, the presence of inter-monomer communities in the active trimer suggests long-distance allosteric coupling between monomers associated with activation (Figure 2). Next, we validated our hypothesis of allosteric activation with an *in vitro* catalytic assay and investigated its implications for inhibitor binding using surface-plasmon resonance (SPR). A similar community analysis was undertaken for the HtrA1 monomer (Supp. Info.).

**Figure 2.**
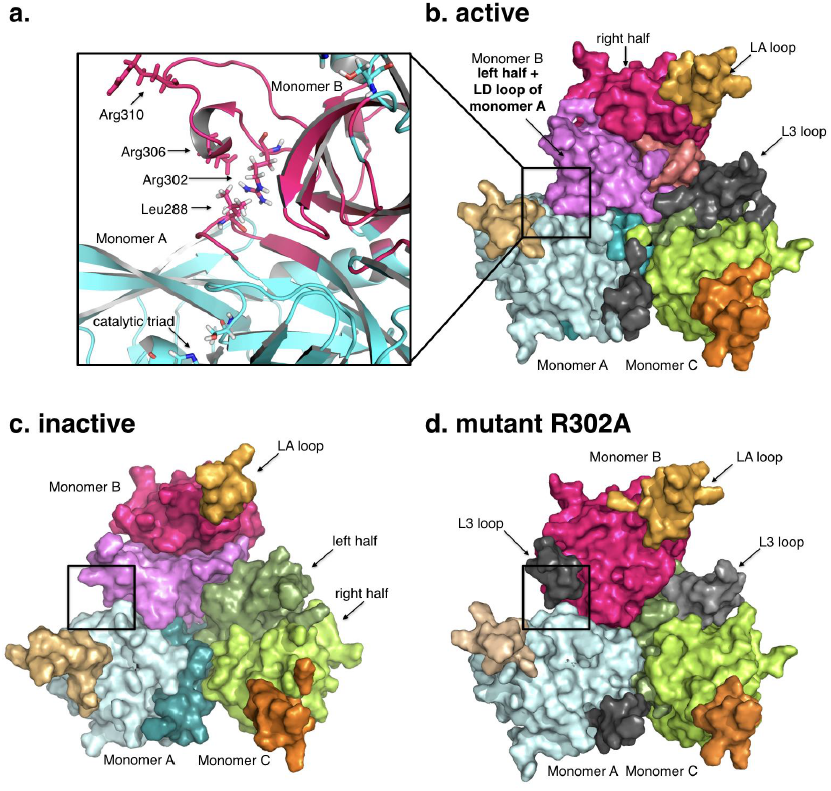
The dynamical communities of the HtrA1 trimeric protein are shown in different colors. (**a**) Intra-monomer community in the active trimer. L3 loop residues (Arg302-Arg310) of monomer B and LD residues (284-290) from monomer A belong to the same dynamic community. (**b**) Active trimer form. (**c**) Inactive trimer form. (**d**) Mutant R302A trimer form. Black regions in (**b**) and (**d**) indicate L3 loop independent communities.

#### HtrA1 trimer dynamic communities

Taken globally, the inactive trimer presents the simplest distribution with almost identical communities in each monomer: two dynamic intra-monomer halves which do not involve inter-monomeric elements, and a third community formed by loop LA (residues 190 to 200) consistent with the inherent flexibility of this region (Figure 2c).

The active trimer, on the other hand, shows a more complex picture. All three monomers also present a basic scheme of two dynamic halves each and a flexible community with the LA loop. However, new dynamic communities arise as specific to the active trimer.(Figure 2b and Supp. Figure S3). Notably, loop L3 (Ser298 to Leu307) appears either independently or coupled to the LD loop located on top of the oxyanion hole of the next monomer (Figure 2a). This dynamic unit bridges adjacent monomers, indicating that in the active form, amino acids in loops L3 and LD from different monomers have correlated motions. Remarkably, the fully inactive trimer does not include such inter-monomer regions.

#### Allosteric activation mechanism

The previous community analysis leads to the hypothesis of an activation mechanism that involves two contiguous monomers. Mechanistically, amino acid side chains in the loop L3 especially the well-conserved Arg302- would play an essential role in HtrA1 activity by interacting with the backbone of the residues Gln289 and Thr291 located at the LD loop of the next monomer. In turn, the LD loop would subsequently interact with the oxyanion forming loop and the catalytic residues, promoting activation. To verify this idea, we first addressed the L3 loop by engineering two variants to disturb the hypothetical allosteric network. Mutation R302A was selected to abolish a possible signal communication between L3 and LD loops, while the double mutant E306A/R310A was designed to substantially modify the L3 loop dynamics, by removing charged residues, without affecting substrate binding. As a proof of concept, first we generated a model for the variant R302A. As expected, the simulation of this variant displayed a loss of the inter-monomeric communities similar to that of the inactive trimer (Figure 2d).

#### In vitro catalytic assays

Following our computational hypothesis, to assess whether inter-monomer communication mediated by the L3 and LD loops is indeed relevant for HtrA1 activity, we measured the catalytic activity of the R302A and E306A/R310A HtrA1 variants *in vitro* (Table 1). As expected from the simulation, R302A presented no catalytic activity while E306A/R310A presented a 2-fold drop in K_cat_. This results confirm that, first, the Arg302 mediated inter-monomeric communication is essential for activity, and second, other mutations in the L3 loop are not perturbatory enough to abolish the activation of the protease (as shown by the catalytic assay).

#### Substrate binding measured with SPR

To investigate the role of the substrate in the activation mechanism and to investigate the effects of R302A on substrate binding, we employed SPR and the peptide-like boronic acid inhibitor DPMFKL-boro-V, which is known to bind to HtrA1 with high affinity^19^. As can be seen in Figure 3a, S328A (inactive control) and R302A variants show a total loss of binding as compared with the wild-type enzyme. This result is compatible with the idea of a stable protease population of inactive conformations, where the Leu345 residue side chain blocks the S1 site. Upon binding of a substrate to an intermediate conformation, if available, the active state would be stabilized, thus assisting in the activation of the remaining trimer units. Consistently, previous studies have already suggested a possible conformational selection mechanism for HtrA1^8^, our evidence now expands this concept to full trimer activation. In this context, in presence of the substrate, the inter-monomer flow of the activation signal is required to stabilize the intermediate active-like conformations that may coexist with the fully inactive trimeric state, as the biochemical experiments with the L3 loop variants (R302A, E306A/R310A) and our simulations suggest.

**Figure 3:**
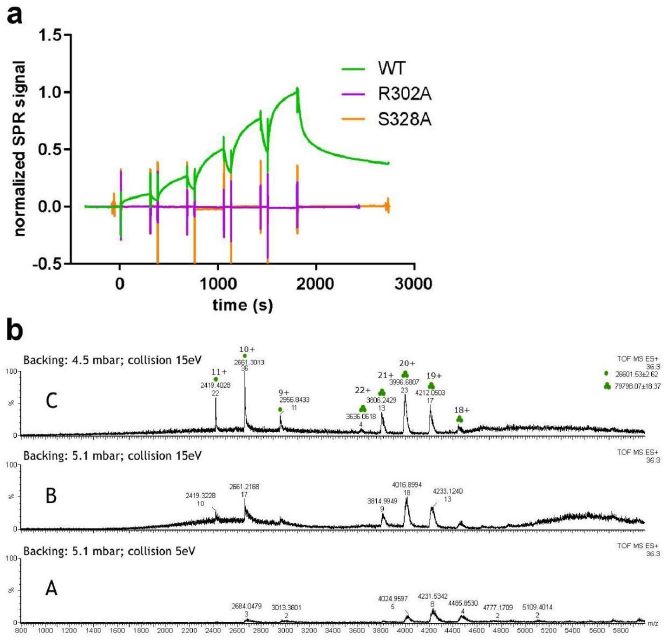
(**a**) Surface plasmon resonance of HtrA1 variants binding to DPMFKL-boro-V: Wild-type HtrA1 catalytic domain (green), mutant R302A (purple) and S328A (dead domain). No binding is observed for R302A or S328A. (**b**) Native mass-spectrometry of HtrA1 mutant R302A. Peaks with three and one green dots correspond to trimer and monomer species respectively. At high-backing pressure (panel **b.A**) mostly trimer species is observed (ratio trimer/monomer = 8/3 → ~2.7) whereas increasing collision energy and/or lowering backing pressure results in an increase in monomeric species (panels **b.B** and **b.C**) and decrease of the ratio *r*. Also in this sample, higher collision energy or backing pressure increase m/z ratio indicating a slight loss of native folded trimeric state. (See Supp. Info. for native MS of wild-type and S328A).

To discard any possible structural disarrangement of the protein due to the introduced mutation that could account for the the lack of binding and proteolytic activity, we verified by native mass spectrometry (MS) that the mutation R302A does not affect the ratio of monomer/trimer populations. Figure 3b shows native MS peaks for constructs wild-type, R302A and the catalytically inactive S328A (negative control) at high and low backing pressure and different collision energies. At high backing pressure, mostly trimer was observed for all constructs, with similar ratios *r* trimer/monomer (WT *r* = ~*6/2* = ~*3*, R302A *r* = ~*8/3* = ~*2.7*, and S328A *r* = ~*20/5* = ~*4.4*). At increasing collision energy and/or lower backing pressure, a decrease in trimeric species is observed in all instances. The rate of decrease in ratio with either backing pressure or collision energy is similar in all cases (For panel C, Δ*r = 0.66* WT, Δ*r = 0.76* R302, Δ*r = 0.6*), indicating that for the mutant R302A, the trimeric species is comparatively stable as the wild type and the negative control S328A (Figure 3b, Figure S6).

#### Role of the LD loop in disease

Having established that perturbations in L3 loop affect both activity and substrate binding *in vitro*, we sought to understand the role of the interfacial LD loop, the L3 loop binding partner, by focusing on naturally-occurring mutations in patients with syndromes linked to HtrA1 dysfunction. Chen et al.^20^ described the clinical phenotype of a CARASIL patient carrying the mutation P285L (LD loop), producing an impaired HtrA1 protease. Similarly, Verdura et al.^6^ identified disease-causing heterozygous mutations S284R, P285Q and F286V in CSVD patients, which produce HtrA1 proteins with residual or decreased catalytic, as measured in an in vitro enzymatic assay of the conditioned supernatants from transfected HEK 293T cells. Still, how heterozygous mutations far from the active site could have such deleterious effect is an open question in that study.

According to the crystallographic evidence, the LD loop is found forming a small helix in the active conformation, while it is disordered in the inactive one. In the helical conformation, the LD loop would be able to interact with the L3 loop of the previous monomer, and with the oxyanion forming loop, promoting a productive arrangement and hence, directly enabling catalysis.

Based on this hypothesis, we ran molecular dynamic simulations for the wild-type and several CSVD-variants to measure the propensity of helicity of the LD loop residues. The results show that the wild-type LD loop is predominantly (95%) helical in the active form (Supp. Table S4, Methods) while all CSVD-related variants present a significant loss of helical propensity (26%-65%). Our simulated model of the variant with disease-causing mutations, therefore, reveals that these mutations affect the secondary structure of the loop LD, disturbing intermonomer communication and precluding activity in complexes which may combine mutant and wild-type HtrA1 forms.

### Significance of inter-monomer allosteric activation in a cellular context

Finally, we assessed the relevance of our findings in a cellular context by overexpressing the full length HtrA1 wild-type and variants, including the PDZ and N-terminal domains in RPE cells. RPE cells are known to overexpress HtrA1 under the pathological condition of AMD, and thus are well-established models for the study of AMD and currently being used in clinical trials^21^. RPE cells were infected by adenoviruses containing the wild-type HtrA1 and the variants S328A (catalytically inactive), R302A and the single mutant R310A.

Western blot analysis of the RPE cell lysates revealed a minor band at 47 kDa, below the intact 54 kDa monomer band, for the wild-type HtrA1 and R310A-carrying vectors (Figure 4a), that agrees with a previously reported self-cleavage band^22^. Instead, the isolating mutation R302A prevents intracellular self-cleavage of the full-length HtrA1 similarly to the negative control S328A, noted by the absence of the corresponding bands (Figure 4a, b).

**Figure 4.**
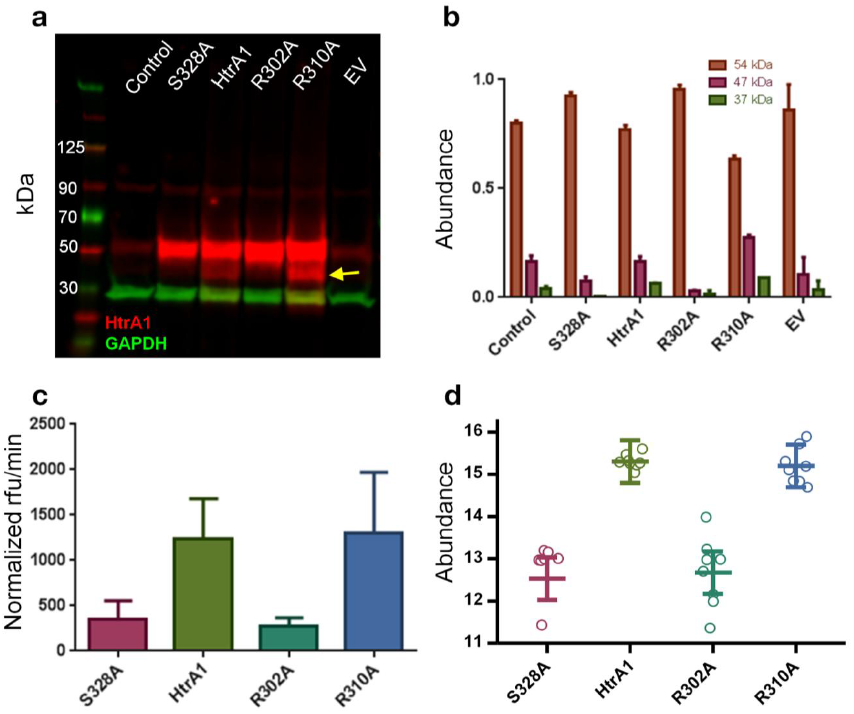
(**a**) Western Blot (WB) of control sample (no virus), and vectors containing: HtrA1 mutant S328A (catalytic inactive control), HtrA1 wild-type, HtrA1 R302A mutant, HtrA1 R310A single mutant and empty vector. The yellow arrow indicates the 47 kDa band corresponding to the HtrA1 self-cleavage product. (**b**) WB quantification normalized to GAPDH (Glyceraldehyde-3-phosphate dehydrogenase). Bands at 37, 47 and 54 kDa correspond to GAPDH, HtrA1 self-cleavage product and HtrA1 monomer respectively. (**c**) HtrA1 in vitro enzymatic assay measured in supernatant from WB, and normalized by total HtrA1 amount (Supp. Figure S5). (**d**) Extracellular HtrA1 activity monitored by the quantification of the semi-tryptic peptide NEQFNWVSR, which is generated through cleavage of clusterin at EQL362-NEQ.

To confirm the inactivity of the R302A variant, we characterized *in vitro* the protein present in the cell supernatant by using an enzymatic assay. Proteolytic activity of the cell supernatant confirms that the enzymatic activity of the variant R302A is also comparable to that of the negative control (Figure 4c). Thus, the R302A mutation has an inhibiting effect but does not impair the secretion of HtrA1 to the extracellular matrix.

On the other hand, the R310A single mutant variant remains similarly active to the wild-type HtrA1 construct, as opposed to the loss of activity shown by the double mutant E306A/R310A in our in vitro biochemical experiments. This suggests that, despite it is located in the L3 loop, R310A alone is not sufficient to perturb the inter monomer communication and prevent activity.

#### Proteomics

Previous studies have demonstrated that HtrA1 is secreted into the extracellular space to digest several extracellular proteins including clusterin^23^. We investigate whether allosteric activation is relevant in the context of HtrA1 extracellular activity. We carry out this analysis by monitoring HtrA1-dependent clusterin cleavage based on a novel cleavage site EQL362-NEQ identified with differential N-terminomic analysis of the secretome of RPE cells. A targeted proteomics assay (selective reaction monitoring, SRM) was developed to quantify the semi-tryptic peptide NEQFNWVSR, which is generated through cleavage of clusterin at EQL362-NEQ by a cellular protease, postulated to be HtrA1 We established and validated the link between the levels of the cleaved clusterin peptide and HtrA1 activity (Supp. Info).

Comparing the levels of the cleaved clusterin peptide allows to characterized HtrA1 activity of our mutants (Figure 4). As expected, the R302A variant presented a similar profile to the negative control S328A: both variants exhibited similar statistically significant decreases in the level of the cleaved clusterin peptide compared to wild-type HtrA1, indicating that these mutations inactivate HtrA1 to similar extents. (Figure 4d, Table 2). HtrA1 levels in the conditioned media of the RPE cells were not perturbed by the R302A mutation in comparison to wild-type (Supp. Figure S5), indicating that the R302A mutation had no effect on HtrA1 secretion. In concordance with the WB data, no significant difference was observed between R310A single mutant and wild-type HtrA1 variants, suggesting that this mutation alone is not sufficient to perturb extracellular HtrA1 activity.

**Table 2:** Statistical significance of the differences (log_2_ fold-change) in the the level of the cleaved clusterin peptide between variants : wild-type, S328A, R310A and S328A. (See the calculation in Supp. Statistics.html).

| Hypothesis | Estimate | p-value |
| --- | --- | --- |
| HtrA1 - EV = 0 | 1.0424 | $2.59 \times 10^{-2}$ |
| S328A - EV = 0 | -1.7369 | $7.65 \times 10^{-5}$ |
| HtrA1 - S328A = 0 | 2.7793 | $1.12 \times 10^{-9}$ |
| R302A - HtrA1 = 0 | 2.6374 | $5.25 \times 10^{-9}$ |
| R310A - HtrA1 = 0 | -0.1093 | $9.96 \times 10^{-1}$ |
| R302A - S328A = 0 | 0.1419 | $9.90 \times 10^{-1}$ |

## DISCUSSION

In human HtrA1, activation has been previously described as a process of allosteric remodelling at the monomer level^11^. In this work, we propose instead an activation mechanism which is triggered by the substrate at the monomer level and cascades globally at the trimer level. This mechanism relies on the communication of the activation signal across contiguous monomers in the HtrA1 trimer. At the monomer level, our combined computational and experimental evidence shows that HtrA1 is self-regulated by a gating mechanism: the enzyme defaults to an inactive state by reversible occlusion of the catalytic site by L2 loop residues. Eventually, binding of the substrate first stabilizes the monomer’s active state, and then the activation signal is allosterically transmitted to other monomers via residues in the L3 loop, and in particular Arg302, facilitating subsequent substrate binding, shown by the wild-type high catalytic efficiency. Furthermore, our SPR experiments show that substrate binding is inhibited in the R302A mutant, showing that binding and inter-monomer communication go hand-in-hand. In this way, HtrA1 would behave as a proficient trimeric machine in which monomers are coordinated and interdependent ratchets that cannot work in isolation.

De Regt et al.^24^ have recently proposed an analogous activation strategy for the bacterial counterparts of HtrA1, based on structural information and mutational analysis of the L3 loop. A triad of residues Thr176, Arg187, Gln200, of which Arg187 is equivalent to Arg302 in HtrA1, mediates inter-monomer allosteric activation of both DegP and also DegS following OMP peptide binding to the PDZ domain. An unsolved question in that study is how OMP-peptide binding to the PDZ domains transmits a signal that results in the remodeling of the protease domain into an active conformation.

Our experimental evidence, based on a computational study of HtrA1 trimer dynamics, shows that this allosteric mechanism, at least in human HtrA1, is possible even in the absence of the PDZ domain and that the L3 and LD loops play an essential role in cascading the activation signal across monomers. Our study establishes that oligomerization is key for the proteolytic activity of HtrA1.

The results of the RPE cellular experiments confirm the relevance of our *in vitro* results, that highlight Arg302 as essential for communicating the allosteric signal across monomers. Perturbation of this cross-monomer signal via the mutation R302A precludes activity both in the presence and in the absence of the PDZ domain. In contrast, a nearby mutation in the L3 loop (R310A), has a negligible effect on activity. Taken as a whole, this indicates that the PDZ domain is not essential for the allosteric activation mechanism to work in HtrA1. Moreover, the isolating R302A mutation does not impede HtrA1 secretion to the extracellular matrix, but can inhibit both extracellular and intracellular HtrA1 protease activity, as measured by the impairment of both clusterin cleavage and HtrA1 self-degradation respectively.

Importantly, our data from native MS shows that the stability of the trimers is not significantly affected by the mutation R302A, confirming that this L3 loop residue is inactivating HtrA1 without disrupting HtrA1 trimeric architecture.

What are the biological implications of such an allosteric activation mechanism in HtrA1? As a protease, HtrA1 should be the subject of tight regulation by the cell in terms of substrate specificity and activity. By default, HtrA1 would be found in an inactive state, maintained by gatekeeper L2 loop residues. Occasionally, the conformational dynamics of loop L2 will partially open access to the catalytic site and subsequent substrate binding would transfer the activation signal to other monomers via L3 and LD loop interaction. Allosteric activation is an elegant strategy for self-regulation, which enables the protease to switch back to an inactive conformation once protein substrates are degraded or activators are removed. Additionally, not all substrates should be equally able to access the HtrA1 enzymatic cavity and activate the enzyme. A concerted strategy of monomer activation, such as the one we propose here, would allow this enzyme to act as a *sensor* that is activated by smaller or ‘easier’ substrates colocalized with more difficult protein targets to be digested. HtrA1 activation would be triggered by the small substrates binding to one monomer, while the other monomers remain active and accessible to other proteins, that by themselves, would not have been able to activate the protease. Such mechanism would also facilitate the degradation of complex tissue and fibrillar proteins such as collagens, fibronectin, fibromodulin, tubulin that have been identified as substrates of HtrA1^25^. Interestingly, in this context, HtrA1 has been found localized in certain brain regions, and attributed a role in the proteolysis of tau and amyloid beta fibrils, as part of HtrA1 house-keeping repertoire^26^.

We have investigated the activation process of the HtrA1 protease by means of computational models, biophysical experiments, proteomics and disease cellular models. Our description of the allosteric regulation of HtrA1 opens up new possibilities for targeting the protease beyond the classical active-site inhibition, and provides an alternative manner of achieving selectivity towards HtrA1. Furthermore, it shows the complementary value of computational molecular dynamics models hand-in-hand with experimental validation at the biochemical, biophysical and cellular levels.

## SIGNIFICANCE

The human protease family HtrA is responsible for preventing protein misfolding and mislocalization, and a key player in several cellular processes. Among these, HtrA1 is implicated in several cancers, and cerebrovascular disease. Our work provides a mechanistic step-by-step description of HtrA1 activation and regulation, which is relevant for drug-targeting this protease. We report that the HtrA1 trimer is regulated by an allosteric mechanism by which monomers relay the activation signal to each other, in a PDZ-domain independent fashion. Our study establishes how HtrA1 oligomerization plays a fundamental role in proteolytic activity. Moreover, it offers a structural explanation for HtrA1-defective pathologies as well as mechanistic insights into the degradation of complex extracellular fibrils such as tubulin, amyloid beta and tau that belong to the repertoire of HtrA1.

## ACKNOWLEDGEMENTS

A.C. and E.M. acknowledge the support of the Roche Postdoctoral Fellowship program. Authors thank Sascha Fauser, Vice President of pRED Ophthalmology, Martin Erkens, Andrew Thomas, Josiane Kohler and Carlos Fenoy. We thank Klaus Müller for inspiration and encouragement. OS acknowledges support from grants DFG (SCHI 871/11-1) and (EXC 294, BIOSS).

### Author contributions

A.C. and P.P performed the computational analysis, planned the experiments and wrote the paper. D.R., E.M., S.H., A.T., F.D., P.J., C.S., C.C. and A.C. carried out the biochemical and biophysical experiments. O.S., R.I., T.D. planned experiments. B.B. performed statistical analysis.

### Competing financial interests

All authors are Hoffman-LaRoche employees or collaborators.

### Data availability

The datasets generated during and/or analysed during the current study are either provided in Supplementary Information or available from the corresponding author on reasonable request.

## METHODS

### Modelling and simulation details

Cartesian coordinates of HtrA1 protease catalytic domain in the active (PDB ID: 3NZI) and inactive (PDB ID: 3NUM,^11^) states were extracted from their respective X-Ray solved structures. Residues were protonated at a pH of 8.0. GROMACS 5.0.5 and the AMBER 03 force field were used to perform all simulation steps^27,28^. To describe the dynamic units of HtrA1 we have used the *MutInf* method described^15,29^ which uses mutual information to identify correlated movements based on simulations in the nanoseconds scale. We have employed the Girvan-Newman algorithm to detect the dynamic communities in our monomeric and trimeric systems. The DSSP program was used to assign the secondary structure to the residues in LD loop (P285 to S287) for each simulation snapshot. Wild-type and variant monomers aggregated simulations, starting from an active conformation, were used by getting snapshots each 5 ns. The percentage of simulation time assigned to a helix was measured by residue. Further computational details are provided in Supp. Information, including Hidden Markov model construction.

### Protein expression

The HtrA1 wild-type catalytic domain (residues Asp161 to Asp369) of the human HtrA1 protease and its R302A, L345G, K346I, E306A/R310A and R310A variants were acquired from Origene. Proteins were expressed in HEK293 cells with a C-terminal FLAG-tag (Myc/DDK) for purification.

### Enzymatic assays

The quenched H2-opt peptide (Mca-IRRVSYSFK(Dnp)K) from Innovagen was used as substrate to carry out all the enzymatic reactions at 25°C^8^.

### Surface plasmon resonance

Proteins were immobilized by standard amino coupling chemistry on a CM5 sensor. Binding of DPMFKL-boro-V was analyzed in duplicates in a titration experiment up to a concentration of 20 µM. Extended details are provided in Supp. Experimental Procedures.

### Native mass-spectrometry

A modified QTOF Ultima (MS-Vision, NL-Almere) mass-spectrometer was used for high-mass measurements. Mass-spectra calibration was done with CsI using the same ionization and declustering conditions. Extended details on sample preparation could be found in Supp. Experimental Procedures.

### Human Fetal RPE Cell Culture and HtrA1 transfection

Cells were purchased from Sciencell (6540), seeded in plastic plates previously coated with Laminin 521 (BioLamina, LN521-03) and cultured for three weeks. We used recombinant adenoviruses containing the human HtrA1 mRNA (GenBank: NM_002775; SIRION Biotech) for the different variants: wild-type, S328A, R302A or R310A.

### Western Blot

analysis was performed similarly as to what is described by Vierkotten et al^30^. Full details in Supp. Info.

### Monitoring HtrA1 activity by targeted proteomics

A targeted proteomics assay was developed using the mass spectrometry-based selected reaction monitoring (SRM) approach for the novel HtrA1-driven clusterin cleavage site EQL362-NEQ. www.msstats.org) in R version 3.3.2 (www.r-project.org)^31^. Peptide intensity values were log_2_ transformed and the median intensities were equalized across all heavy-labeled reference peptides. Peptide summarization to protein abundance was performed using Tukey’s median polish, and the cleaved clusterin peptide NEQFNWVSR was treated as a separate protein. Clusterin cleavage was assessed by adjusting the cleaved peptide abundance for HtrA1 and clusterin content present in the samples, acquired as the residuals of a model containing the two protein amounts. Statistical modeling was performed on these adjusted cleavage values, taking into account the experimental factors and the six different cell types. The following comparisons were made, adjusted for multiple testing: HtrA1 – EV, S328A – EV, HtrA1 – S328A, R302A – HtrA1, R310A – HtrA1, and R302A – S328A. More details can be found in Supporting Statistics.

